# Enhancement of endothelialization by topographical features is mediated by PTP1B-dependent endothelial adherens junctions remodeling

**DOI:** 10.1101/766816

**Authors:** Azita Gorji, Pearlyn Jia Ying Toh, Yi-Chin Toh, Yusuke Toyama, Pakorn Kanchanawong

## Abstract

**Rationale:** Failure of small synthetic vascular grafts is largely due to late endothelialization and has been an ongoing challenge in the treatment of cardiovascular diseases.

**Objective:** Previous strategies developed to promote graft endothelialization include surface topographical modulation and biochemical modifications. However, these have been met with limited success. Importantly, although the integrity of Endothelial Cell (EC) monolayer is crucial for endothelialization, the crosstalk between surface topography and cell-cell connectivity is still not well understood. Here we explored a combined strategy that utilizes both topographical features and pharmacological perturbations.

**Methods and result:** We characterized EC behaviors in response to micron-scale grating topography in conjunction with pharmacological perturbations of endothelial adherens junctions (EAJ) regulators. We studied the EA.hy 926 cell-cell junctions and monolayer integrity using the junctional markers upon the inhibitory effect of EAJ regulator on both planar and grating topographies substrates.

We identified a protein tyrosine phosphatase, PTP1B, as a potent regulator of EAJ stability. Next, we studied the physiologically relevant behaviors of EC using primary human coronary arterial endothelial cells (HCAEC). Our results showed that PTP1B inhibition synergized with grating topographies to modulate EAJ rearrangement, thereby controlling global EC monolayer sheet orientation, connectivity and collective cell migration to promote endothelialization.

Our results showed that PTP1B inhibition synergized with grating topographies to modulate EAJ rearrangement, thereby controlling global EC monolayer sheet orientation, connectivity and collective cell migration and proliferation.

**Conclusion:** The synergistic effect of PTP1B inhibition and grating topographies could be useful for the promotion of endothelialization by enhancing EC migration and proliferation.

## Introduction

Healthy blood vessels are covered by a contiguous monolayer of endothelial cells (ECs), which maintain vasculature hemostasis and serve as a blood-compatible interface. Injured arteries and the associated EC malfunctions result in inflammatory responses, subsequent vessel thickening due to the migration and proliferation of smooth muscle cells, plaque formation, and partial or complete occlusion of the vessels [1]. Such atherosclerotic processes underlie peripheral and cardiovascular diseases, stroke, and myocardial infarction which are among the most prevalent causes of death worldwide [2, 3]. Among the available treatment options, the replacement of the occluded vessel with either an autologous vein or a synthetic vascular graft has been widely practiced. However, while the implantation of large vessels has been a standard procedure, synthetic small vascular graft (diameter <6 mm) has remained problematic. The primary cause of small graft failure has been attributed to the lack of proper endothelialization [4] [5] [6]. Thus, there remains an unmet clinical need for a small vascular graft with higher patency [7]. Strategies employed to improve endothelialization in recent years are typically based on the promotion of EC proliferation and migration of endothelial cells (ECs). These included the pre-seeding of ECs [8], the modification the chemical properties of luminal surface of the grafts, and the use of surface topography [9–11]. However, improvement in endothelialization remained sub-optimal for clinical uses [12, 13], thus calling for further research effort.

Many tissues, particularly the arteries, naturally exhibit topographical features on the micrometer and nanometer scale [14]. Accordingly, many cell types have long been known to be sensitive to nano- and micro-topographical cues [15]. Interestingly, desirable EC behaviors such as EC attachment, alignment, and proliferation are differentially promoted or inhibited by topographical features [16]. For instance, Dolby et al. observed that the attachment and migration of ECs were enhanced on nano- and micro-patterned vascular graft [9], while Murphy and colleagues demonstrated that micron-range grooves decreased the proliferation rates of ECs in the human umbilical vein EC model (HUVECs) [17, 18]. The underlying molecular mechanisms of such EC topography sensing remain poorly understood, however.

The endothelial adherens junctions (EAJs) are adhesive structures mediating the physical connection between adjacent ECs. The EAJs play vital roles in vascular development, monolayer integrity, permeability hemostasis, and vasculature organization [19] [20]. Major EAJ components include VE-cadherin, α-catenin, β-catenin, p120-catenin, N-WASP, Arp2/3, and Vinculin [21]. Multiple signaling pathways have been shown to participate in EAJ regulation. For instance, the binding between VE-cadherin and β-catenin, important for EAJ stability, has been shown to be increased by the activity of Fer tyrosine kinase[22], but reduced by c-Src, Abelson, Fyn, and Pyk2 [23]. Additionally, FAK, Csk and Pak have also been implicated in EAJ regulation [24] [25]. Likewise, numerous tyrosine phosphatases localize to EAJs including PTP1B, SHP2, PTP-μ, DEP-1 and VE-PTP[25]. These play a crucial role in EAJ homeostasis by regulating the phosphorylation of major EAJ constituents [24]. Moreover, EAJs are also profoundly affected by biophysical factors. For example, intracellular tension or external mechanical stimuli (e.g. fluid shear stress) have been shown to positively upregulate the engagement of VE-cadherin-based adhesions to the actin filaments [26] [27] via the mechanosensing functions of VE-cadherin, VEGFR, and PECAM-1[28, 29] [30]. Since previous studies suggested that topographical features alone may be inadequate for the long-term patency of vascular graft [5], an interesting but yet to be explored possibility is that the combination of topographical cues with a pharmacological perturbation of an appropriate molecular target may elicit desirable EC behaviors that facilitate efficient endothelialization. To our knowledge, this approach is yet to be probed in details, in part due to the incomplete understanding of how ECs sense topographical features at the molecular level.

## Methods

### Fabrication of micro- and nano-topographic substrates

Micro- and nano-topographic substrates were fabricated by soft lithography of PDMS (Polydimethylsiloxane) (1), using Sylgard 184 Silicone Elastomer kit (Dow Corning, Michigan, USA). The PDMS polymer was mixed with the curing agent at 1:10 ratio and degassed for 30 min. The photomasks were manufactured by silicon oxide photolithography at the microfabrication core facility, Mechanobiology Institute, Singapore. To generate the substrates, degassed PDMS mixture was deposited onto the silicon oxide molds and degassed again for another 30 min. Subsequently, the PDMS were cured for 2 h in 80 °C oven. The cured PDMS were demolded and cut into the 1×1 cm^2^ pieces to fit inside 24-well tissue culture dishes. Unpatterned PDMS pieces used as the control sample were also fabricated by soft lithography but with the 10 cm tissue culture polystyrene dish (TCPS) as the mold instead. The patterned PDMS pieces were plasma treated for 2 min at 29.6 W power setting (Harrick Plasma, Ithaca, NY, USA), followed by washing in 100% ethanol and UV sterilization for 20 min. The cleaned PDMS pieces were incubated with fibronectin (Sigma-Aldrich) at 1 μg/ml for 1 h prior to the cell seeding. A short degassing was done to promote the adsorption of the fibronectin to the patterned PDMS (2).

### Endothelial Cell Culture

Human endothelial cell lines (EA.hy 926) were obtained from American Type Culture Collection (Manassus, VA), and used with the passage number between 2-10. Cells were verified to be free from mycoplasma infection, and tested monthly. Cells were maintained in Dulbecco’s Modified Eagle’s Medium - high glucose (D7777, Sigma-Aldrich), supplemented with 10% fetal bovine serum (FBS, Gibco) and 1% penicillin-streptomycin (Caisson Biotech, Texas, USA), in 5% CO_2_ incubator. Cell culture media was changed every 2 days. For passaging, cells were detached by incubation with warmed 0.25% trypsin-EDTA (Gibco) for 3 min at 37 °C. Suspended cells were pelleted by centrifugation for 3 min at 700 rpm, and seeded at the ratio of 1:10. HCAEC adapted from Lonza (catalog no. CC-2585) were cultivated in EGM-2mv medium at 37 °C and 5% CO2 and used with the passage number between 5-7. A similar protocol has been flowed for cell passaging.

### Immunofluorescence Microscopy

EA.hy 926 cells were fixed using 4% paraformaldehyde (PFA) in phosphate buffered saline (PBS) for 10 min at room temperature. Subsequently, the cells were permeabilized with 0.1% Triton X-100 for 10 min, followed by 3x washing step in PBS. Fixed samples were blocked by 10% normal goat serum (Invitrogen) for 1 h following by the incubation with the primary antibodies at 4°C overnight or for 1 h at room temperature. Afterward, the samples were washed 3 times in PBS and incubated with the suitable secondary antibodies for 2 h at room temperature. Primary antibodies used are as follows: rabbit anti-YAP, dilution 1:100 (SC-101199, Santa Cruz); mouse anti-β-catenin, dilution 1:200 (610154, BD Transduction Laboratories); mouse anti-vinculin, dilution 1:200 (ab73412, Abcam); mouse anti-phosphotyrosine, dilution 1:200 (05-1050X, Millipore). Cells on PDMS substrates were imaged upside down to be able to achieve higher resolution.

Samples were subjected to imaging using a Nikon A1R laser-scanning confocal microscope equipped with 60X NA1.27 Water Immersion objective lens, image captured by sCMOC (Andor Neo 5.5) camera.

### Drugs and inhibitors

We used 10 μM RK682 to inhibit PTP1B, 10 μM PHPS1 to inhibit SHP2, 10 μM Y-27632 to inhibit ROCK, 50 μM blebbistatin to inhibit Myosin II (All the drugs were purchased from Sigma Aldrich) and DMSO as control. These compounds were incubated with cells for a maximum of 30 mmin. Cells were fixed with 4% PFA immediately after the incubation and without any washing step.

### Gap density quantification

Cells were fixed at 8 h post-seeding using 4% PFA in PBS and stained for β-catenin to label the cell-cell junctions. Samples were imaged using a Nikon A1R confocal microscope with 60X water-immersion objective lens and the images were stitched using the NIS-Elements software to obtain a high-resolution and large field-of-view image. The images were then analyzed using ImageJ to quantify the gap area in each sample.

### Collective and single cell migration assay

The modified scratch assay method (3) was used to generate cell free area on both the planar and patterned PDMS. Briefly, the PDMS substrate was prepared as described above. PDMS pieces were placed inside the 24-well plates. A thin layer of passivated PDMS (3 mm x 10 mm) strip coated with pluronic acid was pasted on top of each PDMS pieces to blocks the cells attachment (Sup figure5). The cell-free areas were genreated perpendicular to the direction of the gratings such that cells could migrate toward the wound area only when the wound is perpendicular to the gratings (3). PDMS samples were plasma treated and coated with fibronectin (1 μg/ml) in PBS for 1 h prior to cell seeding. EA.hy 926 cells were trypsinized and seeded at the density of 300 x 10^3^ per well, and maintained in DMEM growth medium and incubated at 37 °C in 5% CO_2_ incubator before the experiment. After the cells reached 90% confluency, the PDMS strips were removed gently to let the cells migrate toward the cell-free area (wound). The cell nuclei were labeled with NucBlue live ready probes reagent (Invitrogen) to facilitate the tracking of each cell while migrating. Samples were then subjected to imaging for 8 h. For single cell migration assay, the experiment was set up similarly except that the cell density was decreased to 10 x10^3^ per well.

Time-lapse imaging was carried out using Olympus LX81 inverted microscope, equipped with a 10x objective NA 0.30 objective lens and a sCMOS camera (Andor Neo 5.5). Images were acquired for 8 h at 10 min interval. The images were taken in both epifluorescence mode (DAPI channel for the NucBlue-stained nucleus) and phase contrast mode. The DAPI channel images were used for quantification. The mean speed of cell migration and the mean displacement length were determined using the spot tracking function in Imaris v 9.2, ImarisXT (Bitplane Inc.). For each sample, n>200 of tracks were analyzed.

### Image Segmentation and Particle Image Velocimetry Analysis

Image segmentation and Particle Image Velocimetry (PIV) analysis of phase contrast data were performed using in-house software developed in MATLAB. To segment the cell monolayer region, a 25×25 pixels range filter was applied to the raw image. The range-filtered image is a map with high values at large intensity variations region (cell monolayer region) and low values at homogeneous intensity region (wound region). This map was then thresholded and refined by morphological operations to generate a binary mask. The segmented cell monolayer region was then divided into 2 regions of interest based on the distance from the wound edge (i.e., 0-50*μ*m, 50-100*μ*m). To measure instantaneous speed and directional persistence in these regions of interest, Particle Image Velocimetry (PIV) analysis was performed using MatPIV 1.6.1 [73].. Single-pass PIV with a window size of 128×128 pixels and 50% overlap was used. The directional persistence is measured by the cosine of the angle between each velocity vector and the monolayer’s overall moving direction (i.e., from left to right, along the x-axis of the image). The average cosine value ranges from −1 to 1, where a higher value indicates a higher directional similarity between the local cell movement and the overall cell monolayer movement. Note that for data with cells on gratings, pre-processing was performed to remove the gratings on the image before the analysis by subtracting a static background image from the data. The average of images from all time points was calculated to generate the static background image.

### Cell proliferation assay

Cells were seeded with a density of 10×10^3^ cells on both patterned and planar control fibronectin-coated PDMS pieces. At 8 h post seeding cells were treated with PTP1B inhibitor for 1 h followed by EdU staining. Cells were fixed for 10 min with 2% PFA and permeabilized with 0.01% triton X100 for 15 min. Proliferating cells were visualized by EdU staining using Click -iT™ EdU Alexa Fluor^®^ 555 Imaging Kit (Invitrogen, C10338). The fixed samples were imaged using Nikon A1R confocal microscope. The images were then analyzed using Imaris software to count the number of EdU positive nuclei per image area.

### Quantification of YAP Nuclear/Cytoplasmic Ratio

The nuclear/cytoplasmic ratio of YAP was quantified following a previously established method [74]. Immunofluorescence microscopy of YAP in fixed ECs was performed using rabbit anti-YAP, dilution 1:100 (SC-101199, Santa Cruz). Subsequently, the samples were incubated with Alexa Flour 488-conjugated goat anti-rabbit antibody (abcam, 1:500 in 2% BSA) and DAPI (Invitrogen, 100 ng/ml in 2%BSA) to label the nucleus. The samples were then washed 3X with PBS. Imaging was performed using a Nikon A1R laser-scanning confocal microscope equipped with 60X NA1.27 Water Immersion objective lens, and sCMOS camera (Andor Neo 5.5).

The segmentation of cytoplasmic and nuclear region-of-interest (ROI) were performed manually in FIJI (6). Subsequently, the total intensity of the cytoplasmic ROI (*I_total_cell_*), total intensity of the nuclear ROI (*I_total_nuc_*), size of cytoplasm ROI in pixels (*A_cell_*) and size of the nuclear ROI in pixels (*A_nuc_*) were measured. The intensity ratio of nuclear to cytoplasm, *R* was computed as 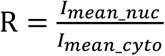, where 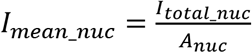 and 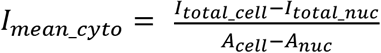 The computation was performed using MATLAB.

### Quantification of Endothelial Adherens Junction Tortuosity

To quantify the tortuosity of EAJs, we used the confocal images of ECs with beta-catenin as the junctional marker, acquired as above. The junctions were traced interactively in FIJI. For each junction, the contour length of the junction, *L*, and the end-to-end distance, *D* were measured. The tortuosity was computed as the ratio of *L/D*. The junction tortuosity value of 1 indicates a straight junction, while large values of the tortuosity correspond to highly convoluted junction topology.

### Statistical analysis

All the experiments were performed in triplicate or greater. Data is presented as mean values and the standard error of mean. Data analysis was carried out using Prism 6.0 (Graphpad Software Inc.). One-way analysis of variance (ANOVA) followed by post-hoc Dunnett’s test was used for multiple comparisons of at least three groups. Unpaired Student’s *t*-test was used for two groups. *P*-values of <0.05 were considered statistically significant.

### Western blotting

Cells were harvested using NE-PER Nuclear and Cytoplasmic Extraction Reagents (ThermoScientific) according to manufacturer’s protocol. The protein concentration of each fraction was measuredusing Pierce bicinchoninic acid (BCA) Protein Assay Kit (ThermoScientific). Protein samples were boiled with 4x NuPAGE LDS Sample Buffer and 10x NuPAGE Sample Reducing Agent (ThermoScientific) for 10 minutes and separated by SDS polyacrylamide gel electrophoresis (SDS-PAGE) on NuPAGE 4-12% Bis-Tris Gel (ThermoScientific) in 1x Bolt MES SDS Running Buffer (Invitrogen) for 60 minutes at 130V.

Separated proteins were transferred to PVDF membrane in 1x FLASHBlot Transfer Buffer (Advansta) for 80 minutes at 90V at 4°C and blocked with 3% BSA for 1h at room temperature. Membranes were incubated with primary antibodies overnight at 4°C with working dilutions stated in Table 1 and washed thrice with TBST for 5 minutes each. Secondary incubation was done at room temperature for 1h and washed again with TBST thrice for 5 minutes.

### Junctional tension measurement by laser ablation

The linear junction in the EA.hy 926 cell monolayers were selectively excised by UV laser nanoscissor ablation performed on a Nikon A1R MP laser scanning confocal microscope, equipped with an ultraviolet laser (PowerChip PNV-0150-100, Team Photonics: 650 nm, 300 ps pulse duration, 1 kHz repetition rate). SIR-Actin, a live-cell actin labeling dye was used to label the junctions. The cut was carried out by using an ImageJ plug-in to control both the actuators and the shutter [43]. Laser ablation was carried out at the z-plane of EAJs using 65 nW laser power focused at the back aperture of the objective lens with an exposure time of 350 ms. Time-lapse confocal images were acquired every 2 s, starting from 3 frames prior to the ablation until 40 frames post-ablation with a scan speed of 1 frame/s and a pinhole size of 74 μm. Image analysis of the recoil speed was performed in ImageJ using the MTrackJ plug-in[44]. This plug-in allows the tracking of the two edges of the cut in subsequent frames and extracts the coordinates of the two points as a function of time. The recoil speed (μm/s) was defined as the rate of change of the two edges[43]. The initial recoil speeds were measured using the data points from the first 2 s after ablation.

## Results

### Microtopography influences morphological features of EAJs

To study the effects of microtopography on the integrity and stability of EC monolayer, we utilized an anisotropic grating pattern with the dimension of 2 μm x 2 μm x 2 μm (spacing x width x depth) (Figure 1A). Such grating dimension was chosen to enhance EC attachment and alignment, based on its similarity to focal adhesion size. To generate the microtopographic substrates, a replica was first fabricated by soft lithography. Subsequently, the patterned substrates were cast using PDMS, following by fibronectin adsorption. Endothelial cells (EA.hy926) were cultured on the patterned substrate, as well as on the planar control, and observed at 8 h post seeding. Since the EAJs are critical structures that regulate EC monolayer integrity, we first compared the EAJ morphology on the grating versus planar substrates, using β-catenin as the immunofluorescence marker for EAJs. As seen in Figure 1B, EAJs adopt two distinct morphologies [31], classified here as the zipper and linear junctions, respectively. The zipper junctions are enriched in VE-cadherin, α-, ϒ-, β-, p120-catenins, and vinculin, with the F-actin fibers aligned perpendicular to the EC border, and with prominent vinculin localization. The zipper junction is also called focal adherens junctions, or cadherin fingers [32] [33], with proposed roles in cell polarity and guidance of collective migration. In contrast, the linear EAJs are distinguished by a smooth appearance, attributed to a local activation of the small GTPase Rac, which reduces tension along the EAJs. In the linear junctions, the F-actin cytoskeleton is oriented parallel to the EAJs [34], with EPLIN as a main linker protein that connect the actin bundle to the VE-cadherin-catenin complex[35].

**Figure 1.**
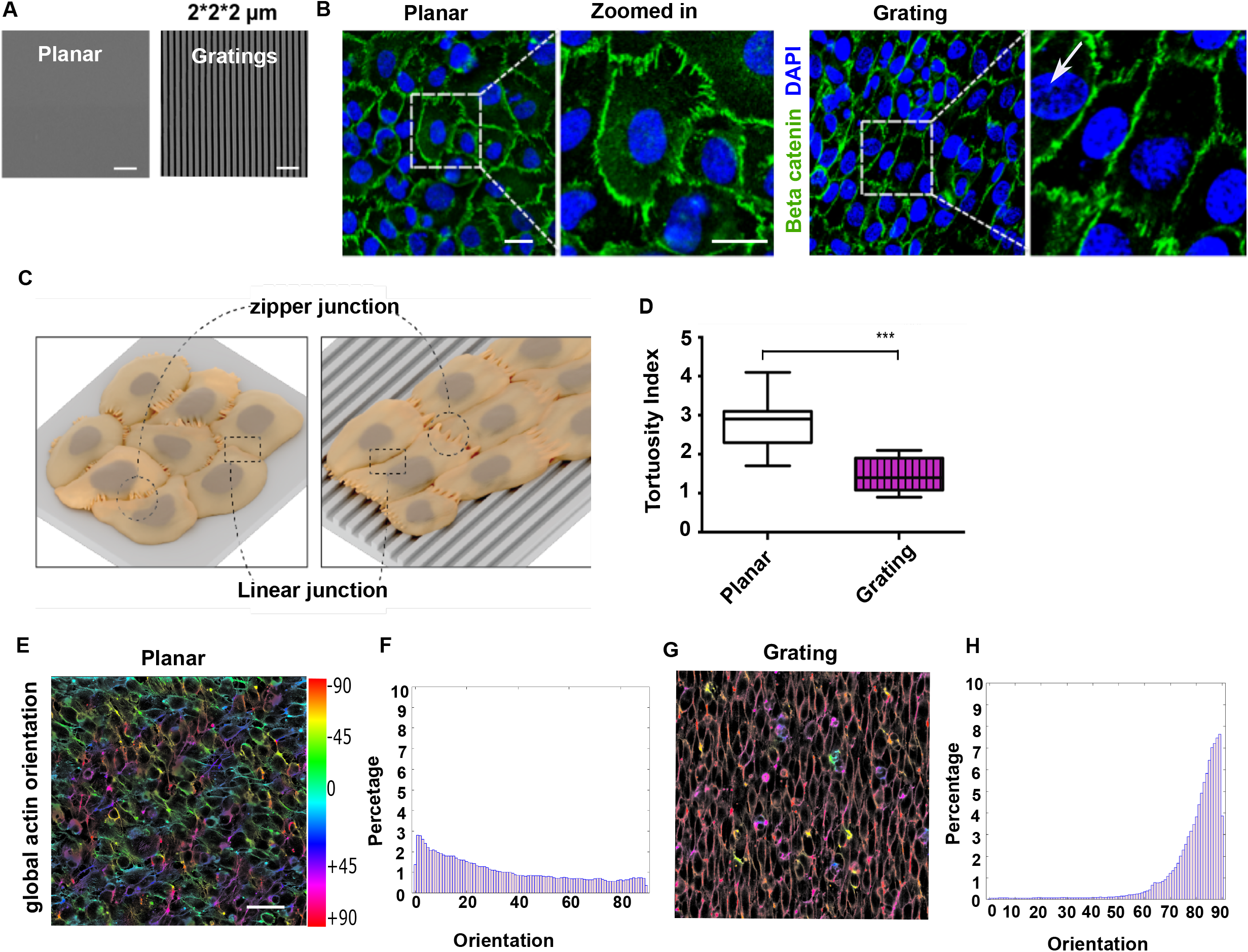
Microtopography influences the morphological features of EAJs. **A**, SEM image of planar and patterned PDMS with grating topographies. Scale bar: 10 μm. **B**, EAJ morphology on planar vs grating substrate. The white arrow indicates grating orientation. Scale bars: 20 μm. **C**, Illustration of EAJs arrangement on planar vs grating substrate. **D**, EAJ morphometric quantification reveals high tortuosity index on planar substrate and low tortuosity index on grating substrate, corresponding to high and low zipper/linear junction ratio, respectively. Data represented as mean±SEM (n =3 to 5). *: p<0.05, **: p <0.005, ***: p <5E-4, ****: p<5E-5. **E-H**, Representative images of the local anisotropy of EC monolayer as quantified from confocal images of F-actin. Color bar, local orientation. **F**, ECs on planar substrate exhibits a largely random orientation of actin fibers. **H**, ECs on grating substrate exhibits a highly anisotropic actin orientation aligned along the grating direction. Scale bars: 50 μm. n = 3-5 independent experiments.

We observed that for ECs on planar substrate, both linear and zipper junctions were formed with no preferential orientation (Figure 1B). In contrast, ECs on gratings are globally aligned along the grating direction. These ECs adopt elongated shapes with relatively smooth EAJs parallel to the grating axis, and “zipper” EAJs along the short edges, perpendicular to the grating axis (Figure 1C). We next used the EAJ morphology as an indicator of endothelial integrity. To quantify the morphology of EAJs, we computed the tortuosity index of EAJs, whereby an index of 1 corresponds to the linear junction, while >1 index value corresponds to a high degree of convolutedness typical of the zipper junctions. The EAJ morphometric analysis revealed that the zipper junctions of ECs on the gratings was largely restricted to the minor axis of the ECs, perpendicular to the gratings orientation, thus accounting for the reduced tortuosity in comparison to cells cultured on planar substrate condition (Figure 1E). Although ECs formed contiguous sheets on both gratings and the planar substrates, we observed significant differences in the global-scale cell organization and cell-scale anisotropy. To quantify global EC orientation, we performed an orientation analysis of the confluent monolayer, using the order parameter calculated from F-actin channel images. On planar substrate, we observed that the actin fibers are oriented without clear directional preference (figure 1F, G). Contrariwise, on gratings a dramatic degree of polarization along the grating axis was observed, as shown in (Figure 1H-I). We concluded that the grating topography promoted global scale orientation that propagated over the entire EC monolayer.

### PTP1B but not SHP2 synergize with topography to stabilize EAJs

Using a candidate-based approach, we next sought to identify biochemical regulators of EAJs that could be pharmacologically perturbed in combination with grating topographies to promote endothelialization-supporting EC phenotypes. The control of tyrosine de/phosphorylation is a major axis for regulating the interaction between VE-cadherin and β-catenin, core proteins of EAJs [36]. Among the tyrosine phosphatases associated with VE-cadherin complex, PTP1B maintains the interaction between VE-cadherin/β-catenin by dephosphorylating Tyr654 residue of β-catenin [37]. Similarly, SHP2 has been shown to induce EAJ reassembly by dephosphorylating β-catenin [38]. Here we used the small molecule inhibitors RK682 and PHPS1 to inhibit the phosphatase activity of PTP1B and SHP2, respectively [39]. As shown in (Figure 2A), we first quantified the effects of PTP1B and SHP2 inhibition on EC morphology. Upon SHP2 inhibition, we found that the aspect ratio of ECs (a metric indicating cell elongation) remained largely unchanged on both planar or grating substrates. Intriguingly, upon PTP1B inhibition, we observed a significant increase in cell elongation on the grating substrate (Figure 2B). The effect of PTP1B inhibition appears to be unique to the grating substrate as ECs on the planar control substrate did not show significant changes in the aspect ratio.

**Figure 2.**
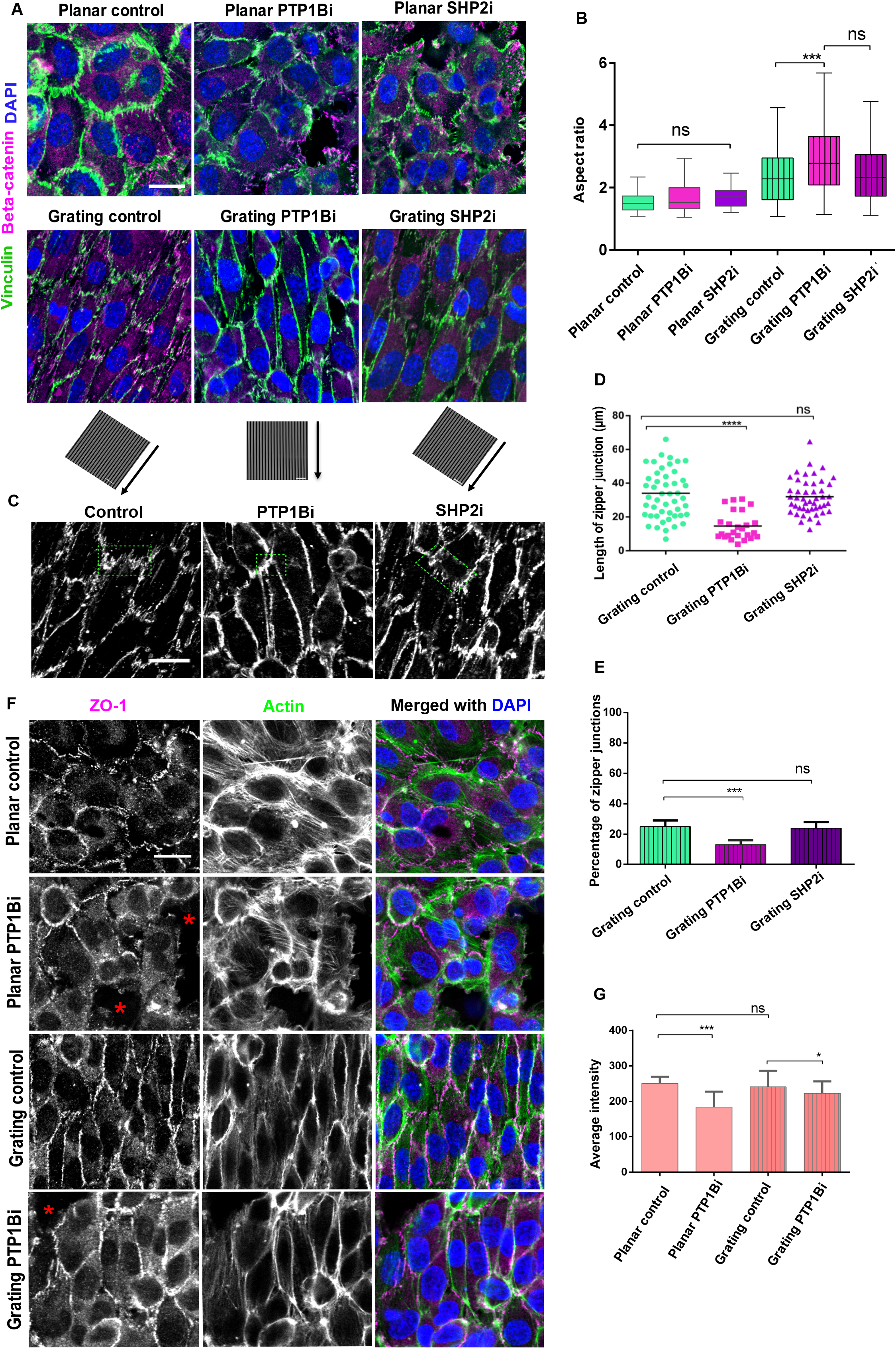
PTP1B, but not SHP2, synergize with grating topography to promote EC elongation. **A**, Confocal images of ECs cultured on planar or grating substrates. Cells were immuno-stained for beta-catenin (green), vinculin (magenta), nucleus (blue). Inhibitor-treated cells were incubated with either PTP1B or SHP2 inhibitors for 30 min before fixation. Grating orientations were indicated on the bottom row. Scale bar: 20 μm. **B**, Aspect ratio of ECs in confluent monolayers. Quantifications were performed in ImageJ (n=10 images). **C**, Zoomed-in views of EAJs comparing before (control) and after pharmacological perturbations. Scale bar: 50 μm **D**, Quantification of zipper junction lengths. **E**, Changes in the percentage of zipper junctions upon pharmacological perturbation. **F**, Merged confocal image of confluent EC monolayer on planar and gratings substrate, immune-stained for ZO-1 (Magenta), F-actin (Green), and DAPI (blue). The red stars indicate the location of gaps in monolayer. Scale bar: 20 μm. G, Quantification of ZO-1 intensity. Significant difference between two groups are indicated as *: p<0.05, **: p <0.005, ***: p <5E-4, ****: p<5E-5, ns: non-significant.

We further characterized the effects of PTP1B inhibition on EAJ morphology of ECs on grating substrate. Using β-catenin as the EAJ marker, our quantification (Figure 2C-D) revealed a two-fold reduction in both the length and number of zipper junctions upon PTP1B inhibition, compared to untreated control. These results implied that PTP1B inhibition may shorten the zipper junctions while increase the formation of linear junctions. Importantly, despite the shortening of the zipper junction due to PTP1B inhibition, we found that key marker such as vinculin still localize to the zipper junctions (Figure S1), suggesting that the molecular composition of the EAJ itself may not be significantly altered.

To evaluate the stability of EAJs under pharmacological perturbations, we quantified the sizes and density of gaps within EC monolayer. ECs were grown on grating topographies to confluency and then treated by PTP1B or SHP2 inhibitors, respectively (Sup figure 1A). From our quantification, we observed that the density of gap increased significantly after the inhibition of SHP2. In particular, the gap density and gap size increased by two-fold upon SHP2 inhibition, in comparison to the control where only small sporadic gaps were observed (indicated by the yellow border in Sup figure 1B). For PTP1B inhibition, we found that the number of small gaps remain largely similar or showing a slight decrease compared to the control sample. Although a small number of large gaps were observed to form, such perturbation appears to not significantly affect the global connectivity of the EC sheet. Taken together, these results suggested that PTP1B inhibition may possess a synergistic effect with the micro-grating topography.

Next, we sought to further assess the integrity of EC monolayer. We probed the level of Zonula Occludens-1 (ZO-1), a component of the tight junction, which served as a major mediator of junctional tension in ECs [40]. ECs were co-immunostained for ZO-1 and β-catenin (yellow stars Figure 2F) and imaged by confocal microscopy. For PTP1B inhibition, we selected regions near the gap for imaging, in order to ascertain whether the gap formation could affect the stability of the adjacent undisrupted junctions. From our quantitative analysis, we found that in control EC monolayer, the intensity of ZO-1 remained largely similar on both planar and grating substrates. For PTP1B inhibition, ECs on planar substrate exhibited a significant reduction in ZO-1 intensity. Interestingly, on the grating substrate the reduction in ZO-1 level is minimal, suggesting that grating topography could help to stabilize EAJs upon PTP1B inhibition.

Subsequently we quantified the level of phosphorylated Tyrosine 731 (pY731) VE-cadherin (Figure S1E) as previous studies have demonstrated that PTP1B dephosphorylated Tyr731 on VE-cadherin, which is associated with the decreased affinity for β-catenin and p120-catenin and EAJ instability[41]. For untreated control ECs, our western blot analysis revealed a decrease in the level of pY731 VE-cadherin for ECs on grating substrate in comparison to planar substrate. These results suggested the regulation of VE-cadherin phosphorylation may comprise one of the underlying mechanisms in the observed synergy of PTP1B inhibition and grating topography in promoting EAJ stability.

We also corroborated these observation by live imaging of ECs on gratings (sup figure 1D). From the analysis of time-lapse movie, we found that cell elongation upon PTP1B inhibition may contribute to the decrease of the monolayer gap. In other words, PTP1B inhibition may enhance the gap closure ability of ECs, In contrast, this effect is not observed upon SHP2 inhibition, thereby accounting for the significantly higher level of gap formation. Altogether these results indicated that PTP1B, but not SHP2, could serve as a suitable pharmacological target for enhancing endothelialization in combination with topographical cues.

### PTP1B-mediated EAJ remodeling is actomyosin-dependent

Given that PTP1B inhibition negatively affected EAJ on planar substrate (Figure 2F), it is surprising that on the grating substrate, PTP1B inhibition promoted EC monolayer integrity. Therefore, we next sought to further characterize EAJ remodelling upon PTP1B inhibition on grating substrate. To investigate the roles of actomyosin contractility, we inhibited ROCK or myosin II using Y-27632 or blebbistatin, respectively, with or without a co-treatment with PTP1B inhibitor. As shown in Figure 3A, we observed a significant disruption of zipper junctions upon the inhibition of either Myosin II or ROCK. Furthermore, from the quantification of cell elongation (Figure 3B), we found that PTP1B-inhibition-induced EC elongation observed earlier (Figure 2B) was largely abolished in ECs co-treated with inhibitors to PTP1B and either ROCK or Myosin II (Figure 3A). Similarly, the inhibition of actomyosin contractility appeared to be associated with a decrease in cell area (Figure 3C). We also investigated the level of phosphorylated myosin light chain (pMLC), which has earlier been shown to correlate with F-actin stress fibers formation in ECs [42]. As shown in (Figure S3A), we observed the co-localizaion of pMLC with the thick actin bundles formed in ECs cultured on the grating substrate. Interestingly, although the total intensity of pMLC staining remained comparable between untreated and PTP1B inhibition, we observed that pMLC may be re-distributed away from cell-cell boundary toward the more centrally-located actin stress fibers (Figure S3B). These results implied that PTP1B inhibition may promote a greater extent of actin stress fiber organization in ECs.

**Figure 3.**
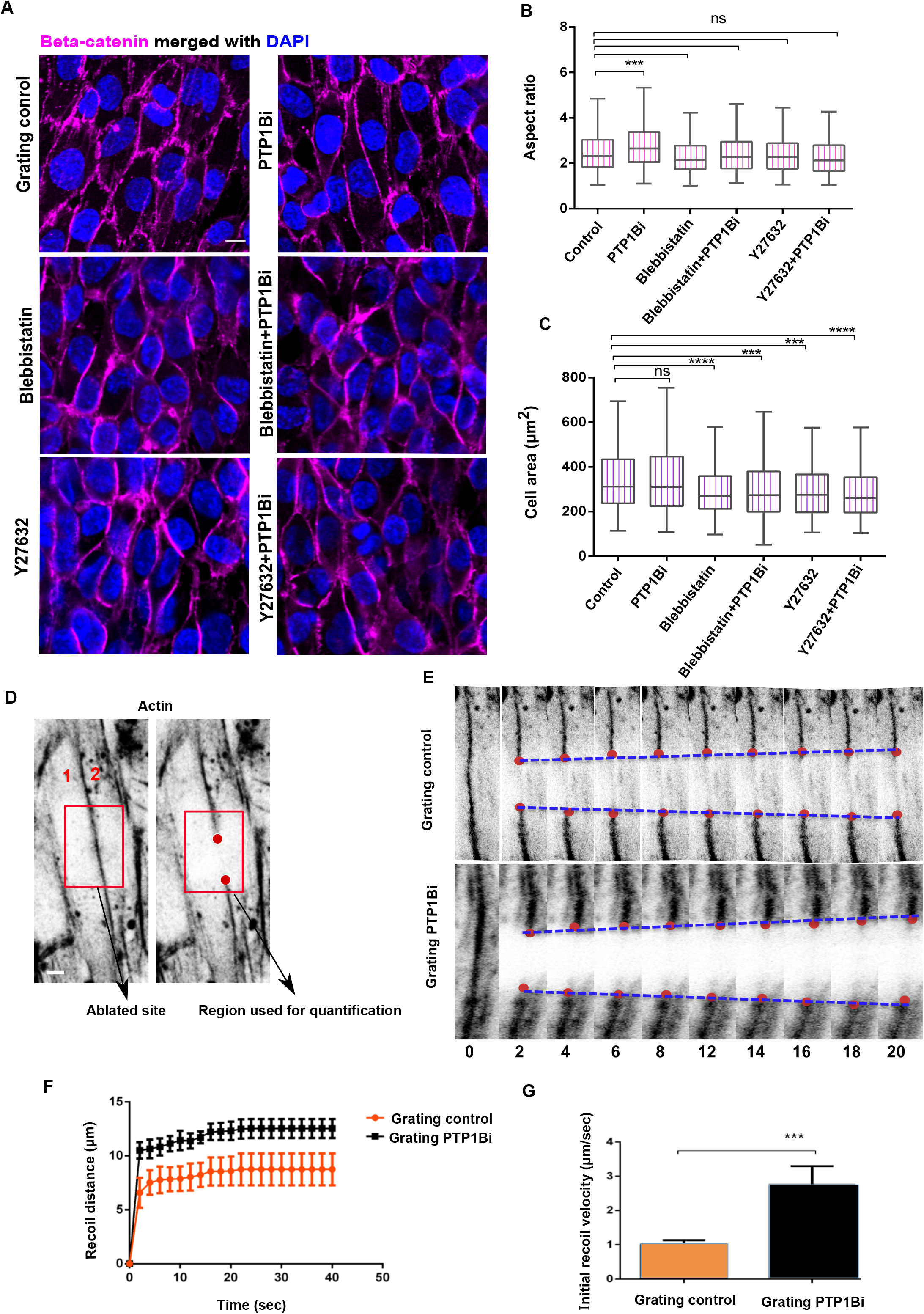
PTP1B-mediated EAJ remodelling is actomyosin-dependent. **A**, Confocal image of confluent EC monolayers on grating substrate, immune-stained for beta-catenin (magenta) and nuclei (blue). Control (untreated) cells were compared with PTP1B inhibition (10 μM RK682), ROCK inhibition (100 μM Y27632), and Myosin II inhibition (50 μM Blebbistatin), or combination thereof. Pharmacological perturbations were performed by 30 min incubation. **B**, Aspect ratio quantification showing that myosin II contractility is necessary for cell elongation after PTP1B inhibition. **C**, Quantification of cell area in response to pharmacological perturbations, n=3-5 independent treatments. **D**, Confocal image of cortical actin at linear EAJ before (left) and after (right) laser ablation. Adjacent cells are marked by 1 or 2. Post-cut junction ends are marked by red dots. **E**, Montage from time-lapse movie of laser ablation experiments, comparing untreated control with PTP1B inhibition. The distances between the edges of recoiling junctions (red dots) were used to quantify the initial recoil velocity. Time interval: 2 s **F**, Junctional recoil distance as a function of time, comparing untreated control with PTP1B inhibition. Data are mean ±SEM form (n= 15). **G**, Initial recoil rate calculated from the initial 2 s post-ablation. Scale bar: 10μm for all images. *: p<0.05, **: p <0.005, ***: p <5E-4, ****: p<5E-5, ns: non-significant.

To more directly probe the effect of PTP1B inhibition on mechanical tension in ECs, we performed laser nanoscissor experiments, which selectively ablate EAJs at specified sites, allowing the EAJ tension to be estimated from the initial recoil rate [43]. ECs were labeled with SIR-actin probe to mark the cell cortex and cell-cell boundary. For our measurements, we chose to ablate the linear junctions which align parallel to the long axis of the ECs, rather than the zipper junctions. We reasoned that tension along the grating axis is likely more relevant for endothelialization. Additionally, since PTP1B inhibition decreased the width of the perpendicular zipper junction, it was more challenging to accurately target the zipper portion of the EAJs (Figure 3D). As shown in Figure 3E, immediately after ablation, the EAJ ruptured and recoiled. The translocation of the ablated junctions was then quantified over time to obtain the recoil velocity rate using the MTrackJ ImageJ plug-in [44], with the initial recoil distance after ablation (L) being depicted as a function of time (Figure 3G). Upon PTP1B inhibition, we observed a significantly faster initial recoil rate in comparison to the untreated control (Figure 3F). This observation thus indicated that upon PTP1B inhibition, EAJ tension was significantly increased along the grating axis. We thus concluded that the effect of PTP1B inhibition on cell elongation and EAJ remodeling could be mediated by actomyosin contractility.

### PTP1B inhibition induces vinculin recruitment to focal adhesions

Given that the interactions between ECs and the substrate topography is mediated by cell-ECM adhesions, while PTP1B inhibition has pronounced effects on EAJ remodeling, we hypothesized that the synergy between PTP1B inhibition and substrate topography may involve the crosstalk between cell-cell and cell-ECM interactions. We focused on vinculin in particular, as it is a major mechanosensitive protein present in both cell-cell and cell-ECM adhesions [45]. Vinculin recruitment is associated with both the tightening of cell-cell junction [46], and reinforcement of cell-ECM focal adhesions [47]. We made use of confocal imaging to compare the localization of endogenous vinculin, as probed by immunofluorescence staining, at the substrate z-section (focal adhesions) and the higher z-section corresponding to EAJ. At the focal adhesions z-plane, we observed a significant increase in vinculin recruitment upon PTP1B inhibition on both grating and planar substrates (Figure 4A and C). In particular, this effect was more pronounced on the grating substrate (Figure 4D). On the other hand, we observed that the vinculin level remained largely similar between the planar control and grating substrates at the EAJ z-plane (Figure 4B and D). Upon PTP1B inhibition, the vinculin expression level was decreased on both planar and grating substrates. Since recent studies have shown that phosphorylated paxillin may be involved in the recruitment of vinculin to focal adhesions [48], we next probed for paxillin in ECs cultured on grating substrate (Figure S4A). Interestingly, we observed a significant increase in the level of paxillin in focal adhesions upon PTP1B inhibition (Figure S4B). In contrast, paxillin was absent from EAJs, consistent with earlier observation [49]. We also probed for Focal Adhesions Kinase (FAK) (Figure S4C). However, the level of FAK appeared to be largely unchanged upon PTP1B inhibition (Figure S4D). Taken together, these results suggested that the effect of the grating topography may primarily involve the strengthening of focal adhesions in a vinculin- and paxillin-dependent fashion.

**Figure 4.**
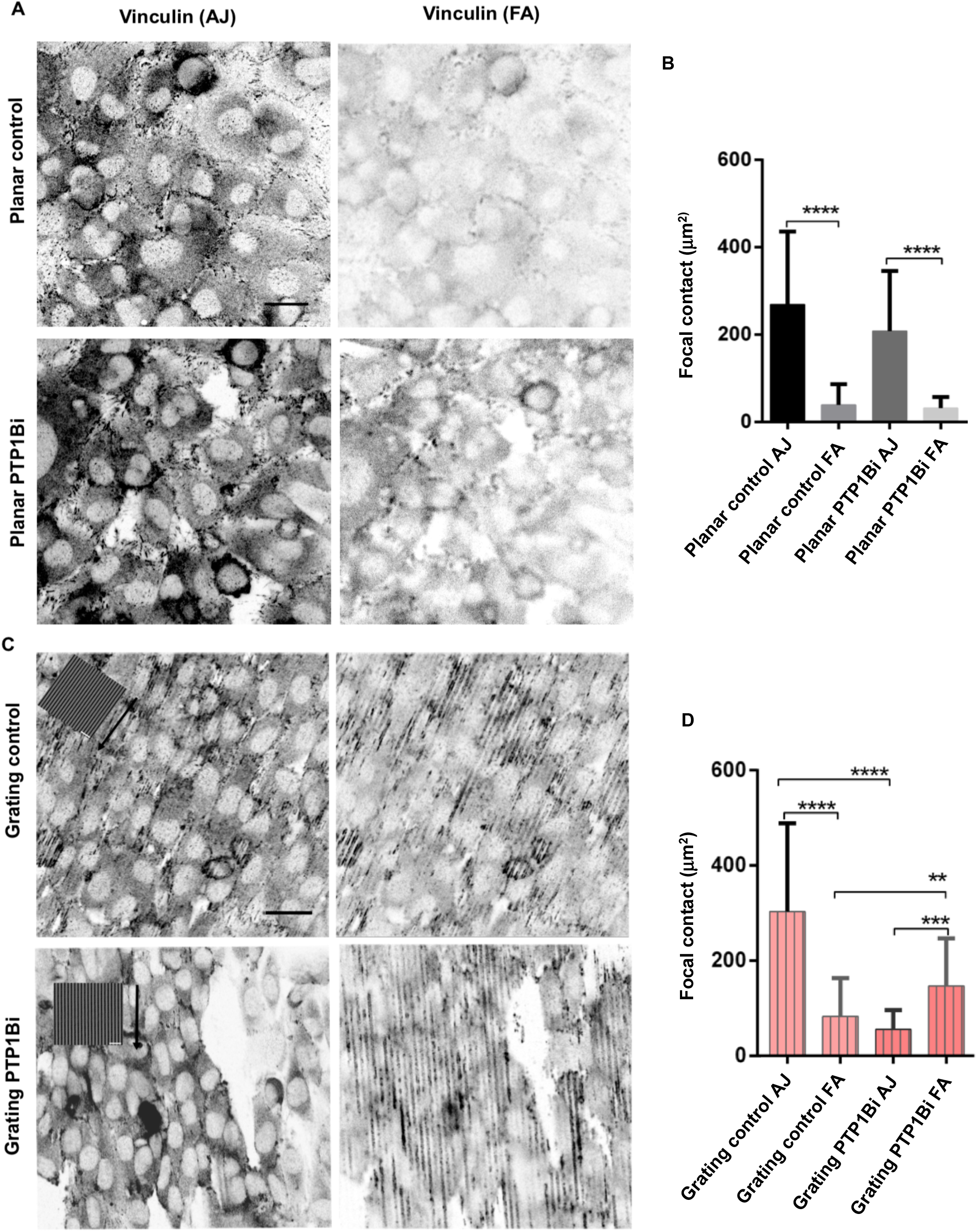
PTP1B inhibition induces vinculin recruitment to focal adhesions. **A** and **C**, Confocal image of EC monolayers under various perturbations on planar (**A**) or grating (**C**) substrate, immune-stained for vinculin, at the EAJ z-plane (left) and focal adhesions (FA) z-plane (right). Grating orientations are shown in insets for C. B and **D**, Quantification of vinculin intensity in ECs on planar (**B**) or grating (**D**) substrate. Vinculin intensity is represented as mean ±SEM per cell, focal contact as μm^2^. n= 10-15 images per condition. Scale bar: 20 μm.

### Grating topography modulates EC motility in synergy with PTP1B inhibition

Anisotropic topographical features are known to promote collective cell migration [50, 51]. Given the importance of EC migration in endothelialization, we next sought to characterize the effect of PTP1B inhibition on EC migration. We monitored the migration of ECs using both primary human coronary artery endothelial cells (HCAEC) (Figure 5A) and the Ea.Hy926 cell line (Figure S5A) to ascertain that the combined effect of PTP1B inhibition and grating typography is common between different EC subtypes. ECs were cultured on PDMS substrates (either planar or grating) to form intact monolayer. Blocks of PDMS were used to maintain a gap region. Gap-closing migration was initiated by removing the PDMS block. Time-lapse movies of ECs after the addition of PTP1B inhibitor were recorded and analyzed by Particle Image Velocimetry (PIV) [52]. For our analysis, we divided the monolayer into the leader section (within 50 μm of the edge), and the follower section (farther than 50 μm from the edge). Our velocity quantification revealed that collective cell migration was significantly accelerated on grating topography (Figure 5B), consistent with previous reports [53, 54]. Furthermore, we found that on grating substrate, PTP1B inhibition increased cell migration velocity further. Such increase in cell migration velocity was registered for both the leader cells (L) and the follower cells (F). In contrast, on planar substrate, collective cell migration was largely unperturbed by PTP1B inhibition (Figure 5B). Here we observed a small increase in the migration velocity of leader cells, but no significant change in the follower cells upon PTP1B inhibition. These results thus further corroborated that the combination of PTP1B inhibition and grating topography resulted in a pro-endothelialization phenotype that included a significant increase in EC collective migration.

**Figure 5.**
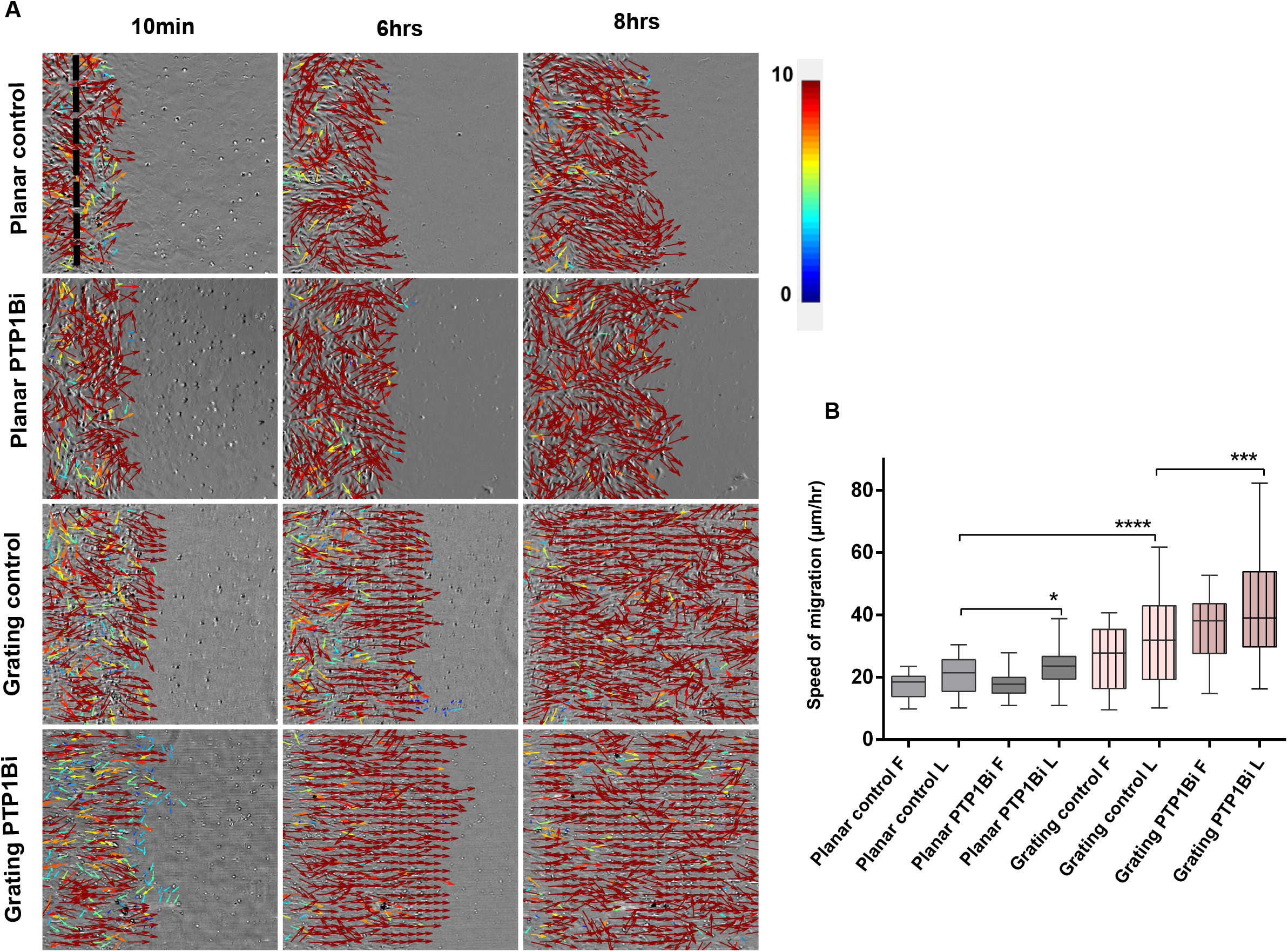
Grating topography modulates EC motility in synergy with PTP1B inhibition. **A**, Montage from PIV analysis of wound-healing migration of HCAEC. Velocity vectors were overlaid on phase-contrast images. Selected frames from 0 h, 3 h, and 8 h are shown. Color bar indicates velocity (μm/h). Speed of migration were analyzed for the leader (L) and follower (F) zones as indicated with black dashed line. **B**, Migration velocity under various perturbations represented as mean±SEM. n=2 movies per condition, experiments were perfomed in triplicate. *: p<0.05, **: p <0.005, ***: p <5E-4, ****: p<5E-5, ns: nonsignificant.

### PTP1B inhibition promotes EC proliferation and YAP nuclear translocation

Since effective endothelialization requires proper proliferation of ECs, we also characterized how PTP1B inhibition may affect EC proliferation on grating substrate. We made use of Edu staining to quantify the percentage of proliferating cells (Figure 6A). HCAECs cultured on grating exhibited a significantly higher proliferation rate upon PTP1B inhibition, in comparison to untreated control (Figure 6B). Since one of the putative molecular links between cell-cell junction stability and proliferation is the mechanosensitive transcription regulator YAP [55], we next sought to characterize how PTP1B inhibition may affect YAP localization. We quantified the nuclear versus cytoplasmic ratio of endogenous YAP localization as probed by immunostaining. We found that PTP1B inhibition significantly promoted the nuclear localization of YAP on both the planar and grating substrate (Figure 6C-D). Thus, although the grating substrate may promote the partial reduction of nuclear YAP, potentially due to the increase in actin cytoskeleton rearrangement [53], PTP1B inhibition appeared to counteract this effect.

**Figure 6.**
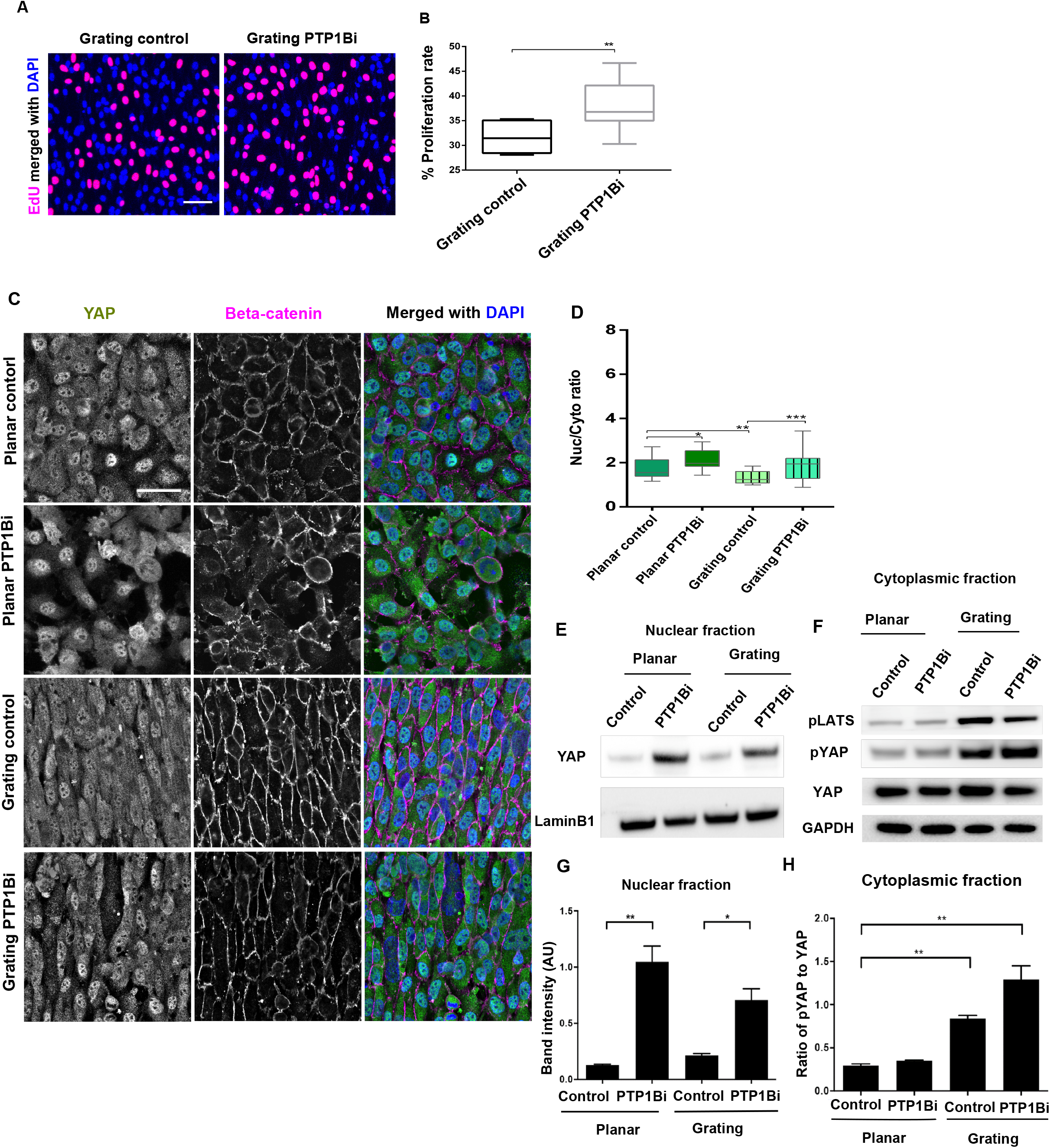
PTP1B inhibition promotes EC proliferation and YAP nuclear translocation. **A**, Confocal image of ECs on grating substrate. EdU staining (Red) is overlaid with DAPI (blue). Scale bar is 50 μm. **B**, Percentage of proliferative cells as quantified from EdU staining. n= 15 images from two independent experiments. **C**, Confocal image of confluent EC monolayers under various perturbations, immune-stained for YAP (Green), beta-catenin (Magenta), and DAPI (Blue). Scale bar is 50 μm. **D**, Quantitative analysis of YAP nucleus/cytoplasmic ratio of YAP. n= 10-20 image for each condition. **E**, Western blot analysis for nuclear **E** and Cytoplasmic **F** fraction of phosphorylated YAP and LATS1/2. **G**, Statistics of nuclear fraction of serine 127-phosphorylated YAP. **H**, Quantification of phosphorylated YAP to YAP in cytoplasm. *: p<0.05, **: p <0.005, ***: p <5E-4, ****: p<5E-5, ns: non-significant.

We next probed the level of phosphorylated and unphosphorylated YAP by Western blotting experiments of either the cytoplasmic or nuclear fractions of ECs. YAP is phosphorylated by active and phosphorylated LATS, preventing its nuclear translocation [56]. Grating topography significantly increased cytoplasmic levels of phosphorylated YAP at S127 (pYAP) as well as phosphorylated LATS1/2, while the amount of total YAP remains the same (Figure 6E). However, increased pYAP did not diminish the levels of nuclear YAP after PTP1B inhibition on grating substrates. Furthermore, in the nuclear fraction, PTP1B inhibition significantly promoted the level of nuclear YAP on both the planar and grating substrates (Figure 6F). These results show that topography increased pLATS activity, therefore downstream pYAP levels and that PTP1B inhibition promoted nuclear translocation of YAP independent of topography. These data implied that PTP1B may be an important upstream regulator of YAP, likely at the EAJs. Further studies will be necessary to dissect the detailed molecular pathways underlying these effects.

## Discussion

In this study, we showed that the combined effects induced by substrate nanotopography and AJ regulation may contribute to promote endothelialization by enhancing EC adhesion, migration and proliferation.

The improvement of small vascular grafts endothelialization has been a long-standing challenge, dating to almost 50 years ago [57]. The initial approach of pre-seeding ECs showed a promising result in reducing the thrombogenicity of the graft, but was hampered by the limited number of cells that can be harvested form the veins, the long-term cell culture prior to seeding and, more importantly, the low retention of the seeded cells[58]. Later on, a number of studies focused on surface chemistry modification to enhance the attachment of ECs. However, achieving an optimal level of EC adhesion, migration and proliferation for endothelial has remained challenging [59, 60]. More recently, it has been increasingly appreciated that the mimicry of the natural topographical features of innate blood vessels can substantially enhance beneficial EC behaviors, with the groove topographies shown to strongly modulate EC adhesion and migration [61]. However, while the maintenance of monolayer integrity is critical for endothelialization, the impact of substrate topography on cell-cell adhesion has not been well characterized. Indeed, to our knowledge, the majority of prior studies tend to overlook cell-cell connectivity and monolayer integrity maintenance, even though these comprise key intermediate steps after EC attachment and before cell proliferation.

In order to fill this gap, in this study we first dissected the role of grating topographies on EC intercellular junctions. Then, using a candidate-based approach, we then focused on SHP2 and PTP1B, major tyrosine protein phosphatases, which serve as important regulators of the AJs. Apart from their regulatory roles at the molecular levels, PTP1B and SHP2 have been identified as potential drug targets for important pathological events related to cardiovascular diseases. For example, PTP1B inhibition has shown great potential in terms of the treatment of obesity [62, 63], type I [64, 65] and II diabetes [66]. In line with this, a recent study also showed that PTP1B deletion in ApoE^-/-^/LysM-PTP1B mice could reduce the size of atherosclerotic plaque [67]. Likewise, SHP2 has been identified as a drug target for acute myeloid leukemia [68, 69] solid tumors [70], and atherosclerosis by negatively regulating smooth muscle cell proliferation [71]. We thereby hypothesize that nanotopography may synergize with the pharmacological inhibition of a specific target to promote desirable EC behaviors amenable to endothelialization.

To dissect such synergestic effect, we focused on 3 major aspects: 1) cell-cell adhesion and monolayer integrity; 2) collective cell migration; and 3) cell proliferation. Our study thus provided key insights into EAJs remodeling during EC collective behaviors.

Using EA.hy926 cells as well as primary human coronary artery cell line (HCAECs) cultured on fibronectin-coated micro-grating and planar PDMS substrates as model surfaces, we showed that grating topographies induced long-range reorganization of EC monolayers as compared to flat surfaces. At the single cell level, cells preferentially aligned along the grating axis leading to well-ordered monolayers at the multicellular scale. This long-range organization is accompanied by specific EAJ arrangement at the level of cell-cell adhesion. We observed distinct morphologies of cell-cell junctions with linear junction formed along the major axis of the cells and zippered junctions along the short axis.

Our study highlighted the roles of the cell-cell junctions as the foremost regulator of monolayer integrity. Upon PTP1B or SHP2 inhibitions, differential responses of ECs were observed. SHP2 inhibition leads to the formation of gaps within the monolayers but no significant alteration in EC morphology (Figure Sup 1B). On the other hand, upon PTP1B inhibition, ECs exhibited a higher aspect ratio and became more contractile, while still retaining their junctional stability, forming more prominent focal adhesions at the cell-substrate interface through the additional recruitment of vinculin and paxillin.

Furthermore, our data suggest that the combined effects of PTP1B inhibition and grating topographies favor collective cell migration by increasing the mobility of both leader and follower cells in the group. We also confirmed the positive effect of PTP1B inhibitor on cell proliferation on gratings as ilustrated in model of our study in (Figure7A). We note that YAP activation has been documented as a positive regulator of cell proliferation in diverse cell types with a recent study showing that that VE-cadherin-PI3/Akt pathway may directly regulates YAP activation and nuclear translocation to modulate of angioprotein [72]. According to the current knowledge regarding YAP as regulator of both EAJs and proliferation, we further investigated how PTP1B could affect YAP regulation through EAJ stability. By immunofluorescent staining of YAP and its phosphorylated level, we found that PTP1B inhibition induced YAP nuclear translocation independent of grating topographies (Figure 7B).

**Figure 7.**
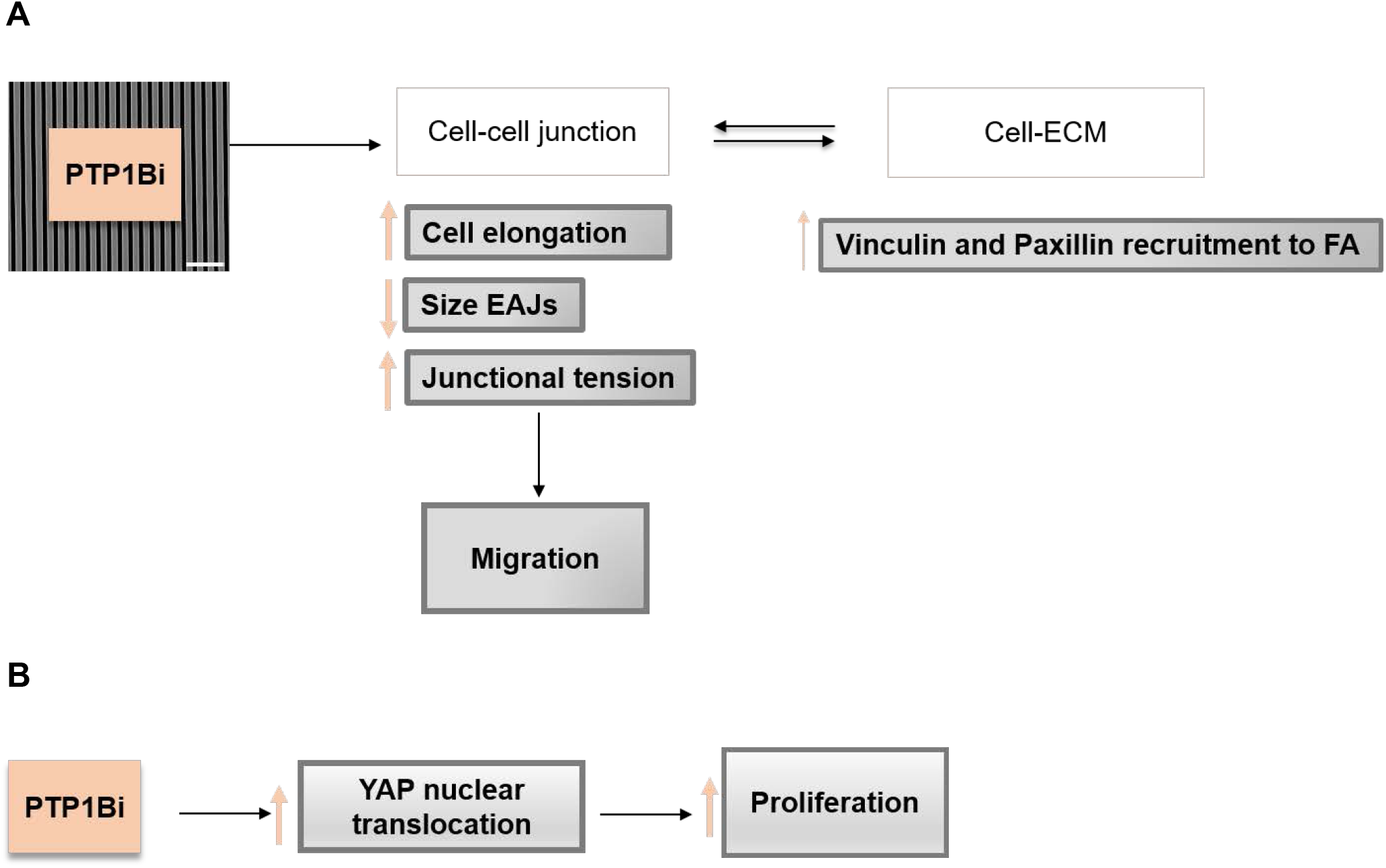
Proposed model for the synergistic effect of PTP1B inhibitor and grating topographies on migration and proliferation through EAJs remodeling. **A**, The synergistic effect of PTP1B inhibition with the topographical features at the cell-cell junction site of EC lead to cell elongation along the grating axis, EAJ compaction, and an increase in junctional tension. Subsequently, the crosstalk between the cell-cell junction and cell-extracellular matrix adhesion promotes Vinculin and Paxilin recruitment to the focal adhesion. **B**, PTP1Bi could result in YAP nuclear translocation independent of the gratings. Globally, EAJs remodeling induced by grating and PTP1Bi, together with YAP activation, promote collective cell migration and proliferation that may enhance endothelialization.

Altogether, our study introduces a promising avenue to improve endothelialization by combining the biomaterial approach of promoting the physical coupling of cells to their underlying substrate with the pharmacological approach of fine-tuning the regulation of AJs. We demonstrated the synergistic effect of PTP1B inhibitor with the grating topographies in promoting beneficial EC phenotypes for endothelialization. We anticipate that such combined approach could not only provide a better understanding of how the EC monolayers respond to physical and biochemical cues but also open new routes towards the improved endothelialization of vascular grafts.

## Acknowledgements

We gratefully acknowledge finanical support from the Mechanobiology Institute (MBI) seed funding (to P. K.), MBI graduate scholarship (to A.G.), and the Ministry of Education Academic Research Fund Tier 2 (MOE-T2-1-124, to P.K.). We thank Hui Ting Ong for assistance with image analysis.

