## Supplementary figures for "Enhancement of endothelialization by topographical features is mediated by PTP1B-dependent endothelial adherens junctions remodeling"

A

beta-catenin Actin Vinculin Merged

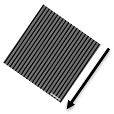

Grating control

Grating PTP1Bi

B

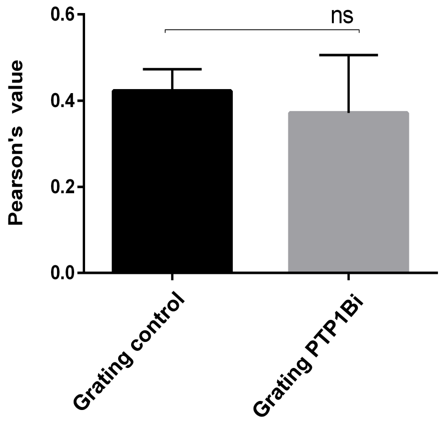

**Sup figure 1. Vinculin remains in EAJ after PTP1B inhibition**

**A**, Immunofluorescence images of EC monolayer on grating substrate, stained for beta-catenin (green), actin (blue), and vinculin (red). PTP1B inhibitor treatment was for a duration of 30 min. Scale bar: 20μm. **B, Co**-localization analysis of vinculin and beta-catenin.

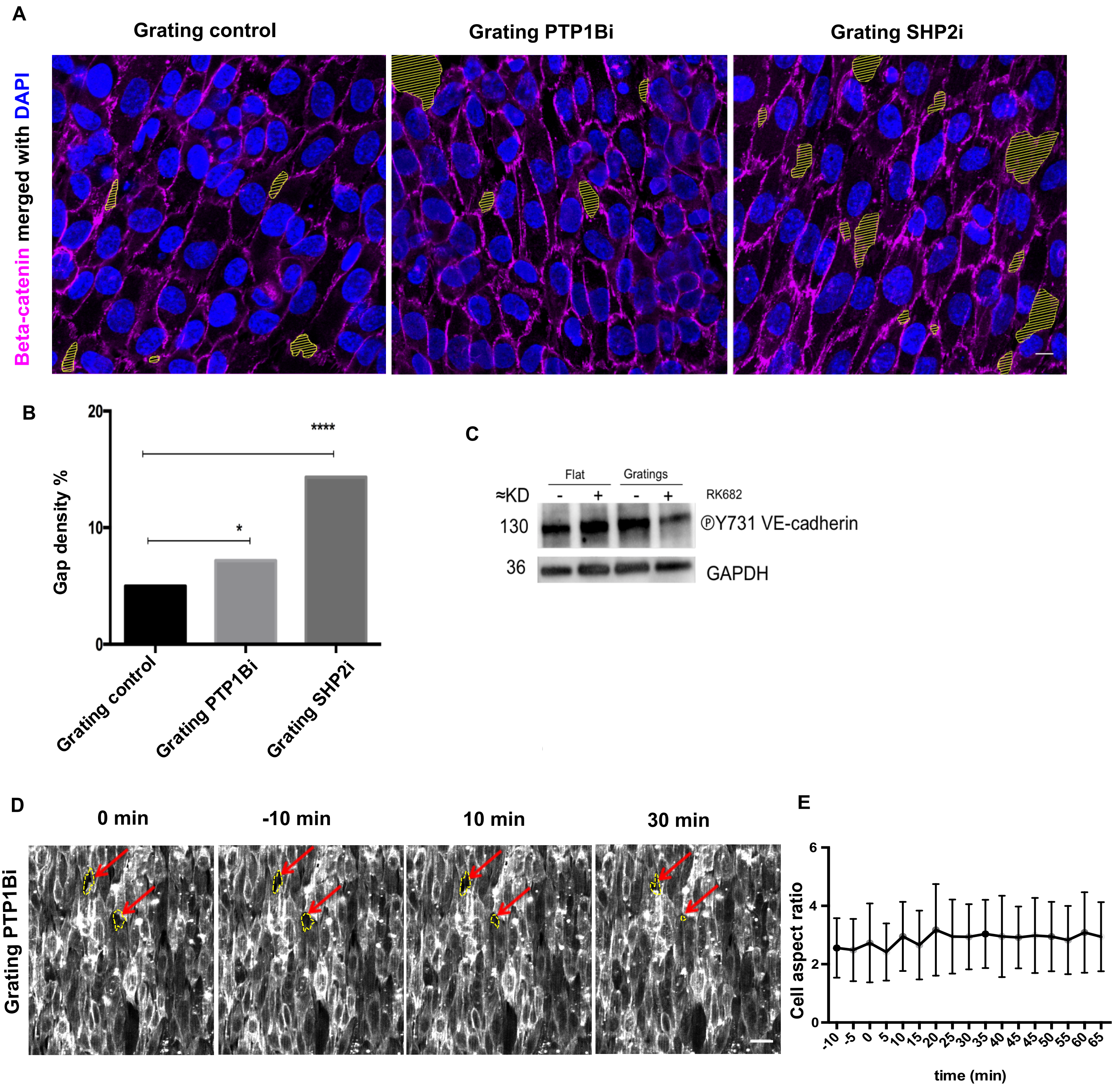
A
Grating control
Grating PTP1Bi
Grating SHP2i
Beta-catenin merged with DAPI
B
Gap density %
****
*
Grating control
Grating PTP1Bi
Grating SHP2i
C
Flat
Gratings
≈KD
RK682
130
36
ⓅY731 VE-cadherin
GAPDH
D
0 min
-10 min
10 min
30 min
Grating PTP1Bi
E
Cell aspect ratio
time (min)

**Sup figure 2. PTP1B inhibition reduce monolayer gap size by promoting cell elongation**
**A**, Confocal image of confluent EC monolayer treated with PTP1B or SHP2 inhibitor, immunostained for beta-catenin (magenta) and nucleus (blue).Yellow marking indicates intra-monolayer gaps. Gaps were quantified after 30 mins of treatment. . n= 5-8 treatment. **B**, Quantitative analysis of gap density (percentage of total monolayer area). **C**, Western blot analysis of phosphorylated Tyrosine 731 of VE-cadherin in ECs under various perturbations, in comparison to GAPDH loading control. Results are representative of three experiments.
**D**, Montage of selected frames from the time-lapse movies showing the response of EC monolayer to PTP1B inhibition (10 μM RK682). Live-cell probes for actin (SIR-actin) was used to label F-actin. Time indicated is relative to PTP1B inhibition addition (t = 0). **E**, Time-course of cell elongation as quantified from time-lapse movies. Scale bar: 10μm.

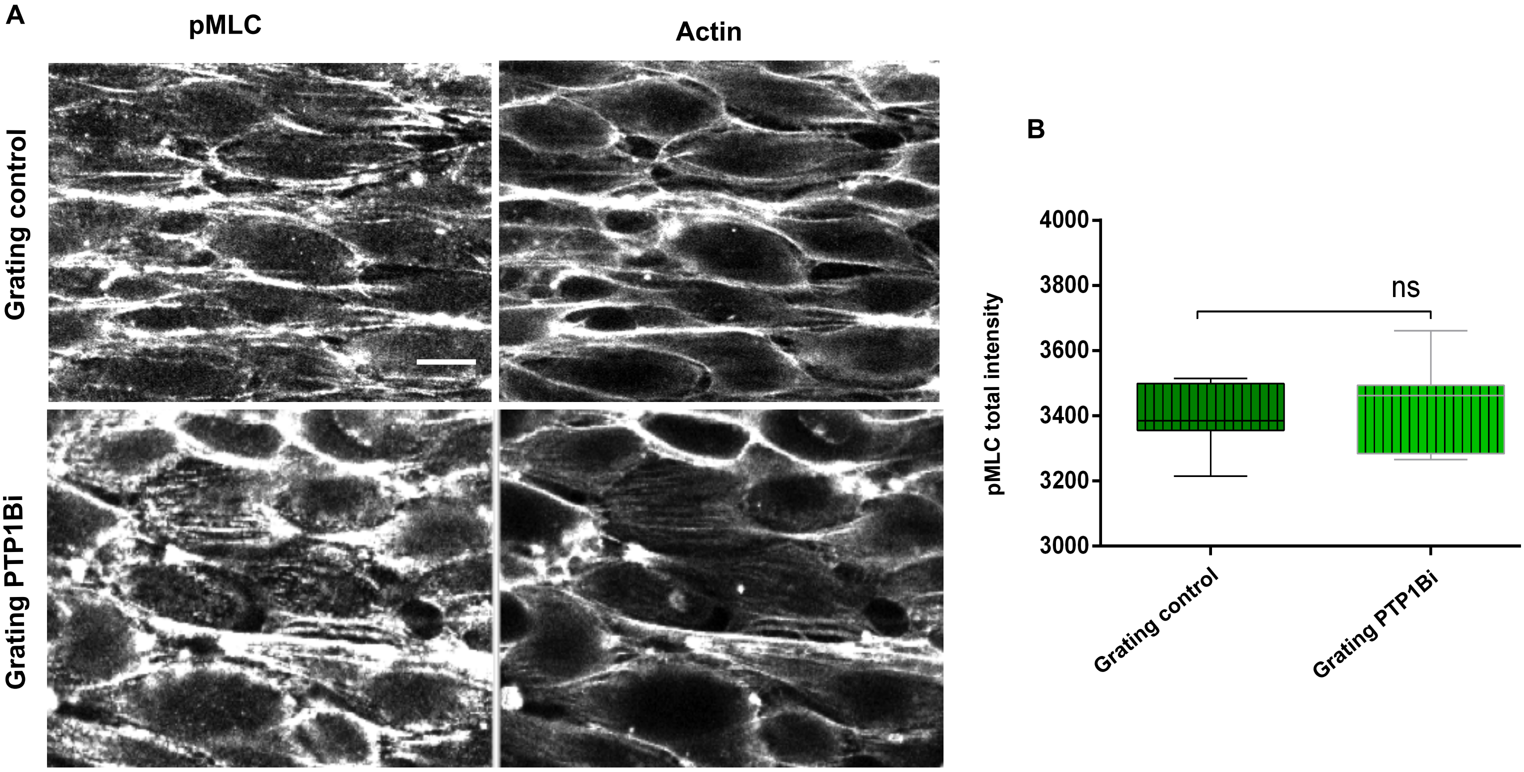
A
pMLC
Actin
Grating control
Grating PTP1Bi
B
4000
3800
3600
3400
3200
3000
pMLC total intensity
ns
Grating control
Grating PTP1Bi

**Sup figure 3.The effect of PTP1B inhibition on the spatial localization of phosphorylated MLC**.

**A**, Confocal image of ECs on grating substrate immunostained for phosphorylated MLC (Magenta) and actin (Green). Scale bar: 50 μm.**B**, Quantitative analysis of pMLC intensity for control (upper panel) and PTP1B inhibition (lower panel). n=10-15 images from two independent experiment. *$P < 0.05$, **$P < 0.01$,***$P < 0.001$, NS non-significant.

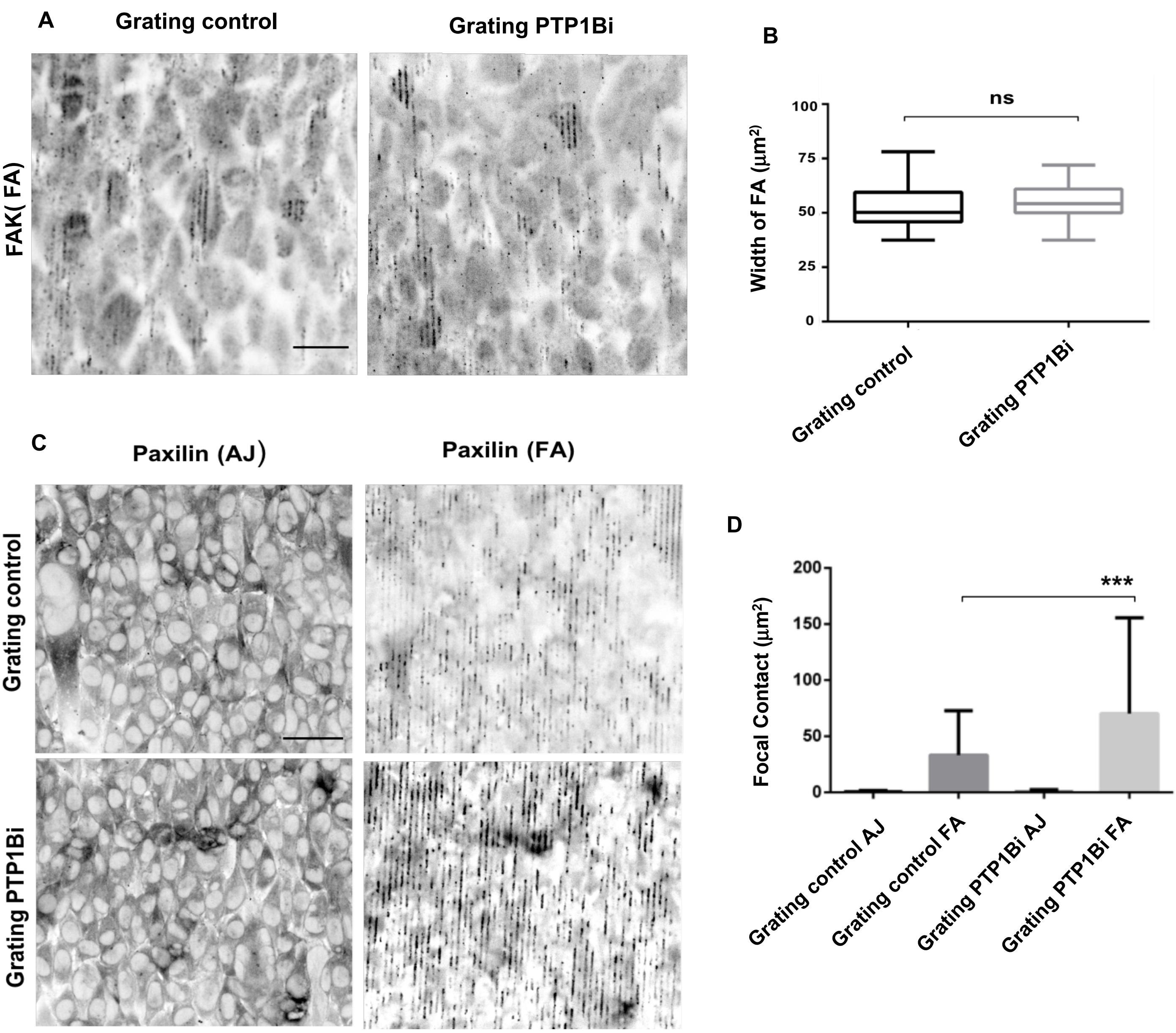

A
Grating control
Grating PTP1Bi
FAK( FA)
B
ns
Width of FA (μm²)
100
75
50
25
0
Grating control
Grating PTP1Bi
C
Paxilin (AJ)
Paxilin (FA)
Grating control
Grating PTP1Bi
D
***
Focal Contact (μm²)
200
150
100
50
0
Grating control AJ
Grating control FA
Grating PTP1Bi AJ
Grating PTP1Bi FA

**Sup figure 4. Paxillin, but not FAK, is recruited to focal adhesions of ECs on grating upon PTP1B inhibition**

**A**, Confocal images of ECs on grating substrates, immunostained for FAK, and imaged at the focal adhesions (FA) z-plane. **B,** Quantitative analysis of FAK intensity at FA z-plane. **C**, Confocal images of ECs on grating substrates, immunostained for paxillin, and imaged at the EAJ z-plane (AJ) or focal adhesions z-plane (FA). **D**, Quantification of paxillin intensity at AJ and FA z-planes. Scale bars: 50µm. $p< 0.05$,$^{**}P < 0.01$, $^{***}P < 0.001$, NS non-significant.

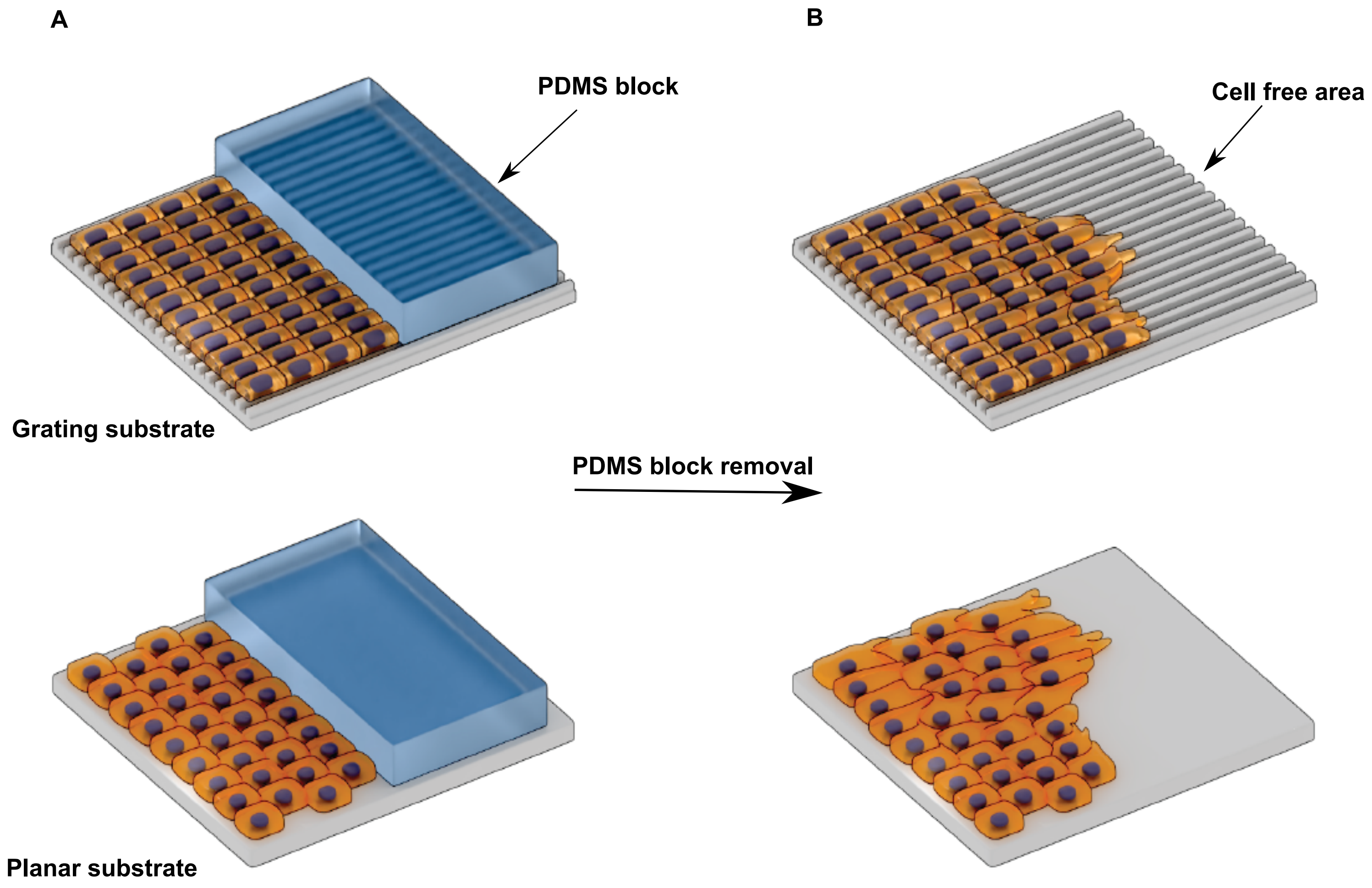
A
B
PDMS block
Cell free area
Grating substrate
PDMS block removal
Planar substrate

**Sup Figure 5. Schematic represents the in vitro scratch assay using PDMS blocks. A**, Cells seeded on Grating and planar fibronectin coated PDMS substrates, the PDMS block used to inhibit cell attachment **B**, Cell migrating towards the cell free area after PDMS block removal.

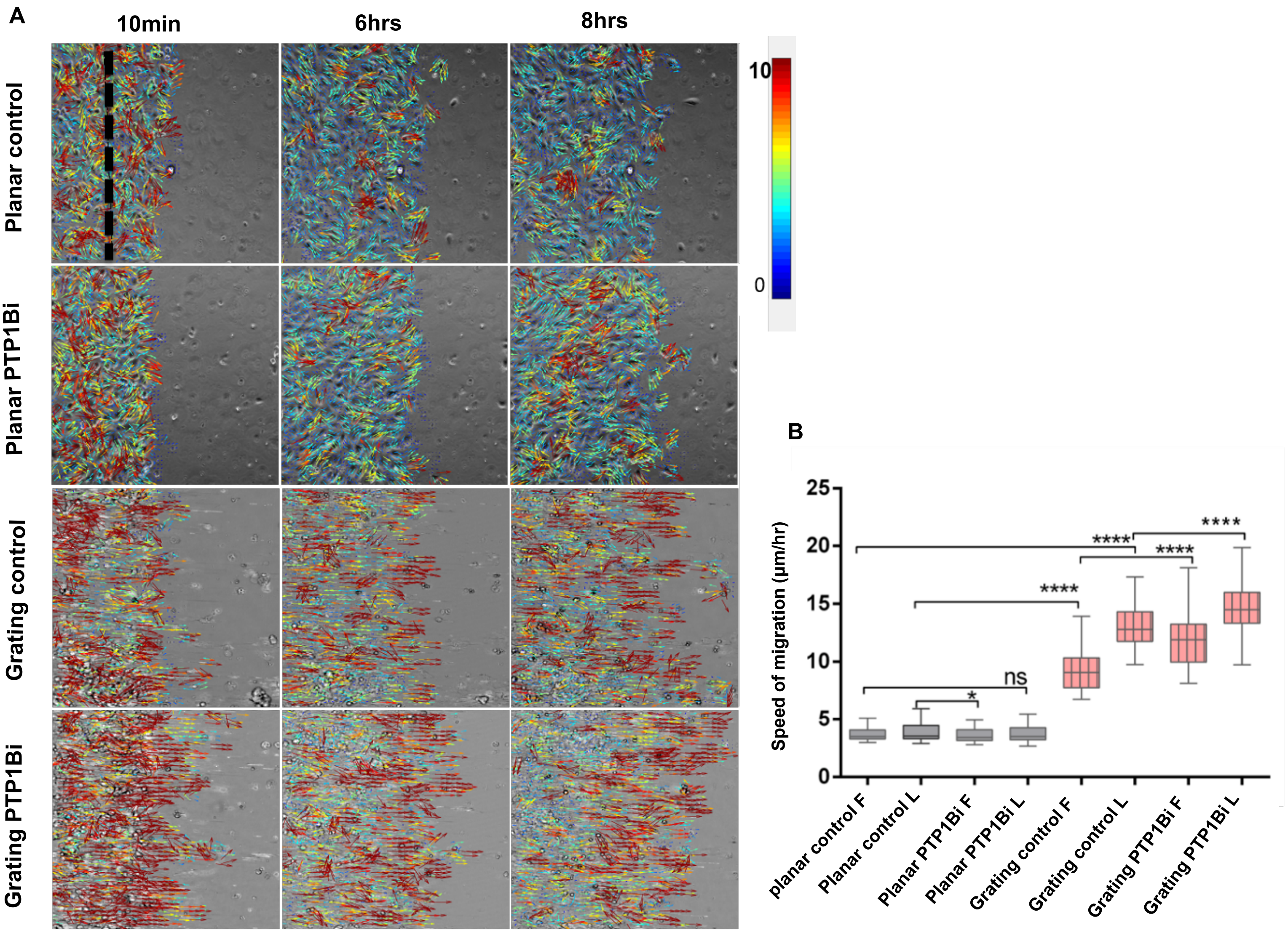
A
10min
6hrs
8hrs
Planar control
Planar PTP1Bi
Grating control
Grating PTP1Bi
10
0
B
Speed of migration (µm/hr)
25
20
15
10
5
0
****
****
****
****
ns
*
planar control F
Planar control L
Planar PTP1Bi F
Planar PTP1Bi L
Grating control F
Grating control L
Grating PTP1Bi F
Grating PTP1Bi L

**Sup figure 6**. **Grating topography modulates EA.hy 926 cell's migration speed in synergy with PTP1B inhibition A**, Montage from PIV analysis of wound-healing migration of EAhy 926. Velocity were analyzed for the leader (L) and follower (F) zones as indicated with black dashed line. **B**, Migration velocity under various perturbations represented as mean ±SEM. n=3 movies per condition, experiments were perfomed in triplicate. *: p<0.05, **: p <0.005, ***: p <5E-4, ****: p<5E-5, NS non-significant.
